# Increased Brain Volumetric Measurement Precision from Multi-Site 3D T1-weighted 3T Magnetic Resonance Imaging by Correcting Geometric Distortions

**DOI:** 10.1101/2021.11.29.469919

**Authors:** Nuwan D. Nanayakkara, Stephen R. Arnott, Christopher J.M. Scott, Igor Solovey, Shuai Liang, Vladimir S. Fonov, Tom Gee, Dana N. Broberg, Seyyed M.H. Haddad, Joel Ramirez, Courtney Berezuk, Melissa Holmes, Sabrina Adamo, Miracle Ozzoude, Athena Theyers, Sujeevini Sujanthan, Mojdeh Zamyadi, Leanne Casaubon, Dar Dowlatshahi, Jennifer Mandzia, Demetrios Sahlas, Gustavo Saposnik, Ayman Hassan, Richard H. Swartz, Stephen C. Strother, Gregory M. Szilagyi, Sandra E. Black, Sean Symons, ONDRI Investigators, Robert Bartha

## Abstract

Magnetic resonance imaging (MRI) scanner-specific geometric distortions may contribute to scanner induced variability and decrease volumetric measurement precision for multi-site studies. The purpose of this study was to determine whether geometric distortion correction increases the precision of brain volumetric measurements in a multi-site multi-scanner study. Geometric distortion variation was quantified over a one-year period at 10 sites using the distortion fields estimated from monthly 3D T1-weighted MRI geometrical phantom scans. The variability of volume and distance measurements were quantified using synthetic volumes and a standard quantitative MRI (qMRI) phantom. The effects of geometric distortion corrections on MRI derived volumetric measurements of the human brain were assessed in two subjects scanned on each of the 10 MRI scanners and in 150 subjects with cerebrovascaular disease (CVD) acquired across imaging sites.

Geometric distortions were found to vary substantially between different MRI scanners but were relatively stable on each scanner over a one-year interval. Geometric distortions varied spatially, increasing in severity with distance from the magnet isocenter. In measurements made with the qMRI phantom, the geometric distortion correction decreased the standard deviation of volumetric assessments by 35% and distance measurements by 42%. The average coefficient of variance decreased by 16% in gray matter and white matter volume estimates in the two subjects scanned on the 10 MRI scanners. Geometric distortion correction using an up-to-date correction field is recommended to increase precision in volumetric measurements made from MRI images.

## Introduction

Degenerative brain diseases are amongst the most prominent and costly (1) disorders affecting the ageing population. These disorders typically change or destroy personality, memories, or cognitive abilities, and are ultimately fatal. The development of imaging biomarkers for the early identification and management of neurodegenerative diseases increasingly involves large scale studies combining magnetic resonance imaging (MRI) data acquired at multiple sites (2–5). The Ontario Neurodegenerative Disease Research Initiative (ONDRI) is a province-wide study that recruited more than 550 subjects from disease groups including amyotrophic lateral sclerosis, fronto-temporal dementia, Parkinson’s disease, cerebrovascular disease, Alzheimer’s disease and mild cognitive impairment (4). The ONDRI was a prospective cohort study to characterize the contribution of vascular disease in neurodegenerative conditions using indicators including MRI measures of vascular disease burden, gait impairment, detailed neuropsychological testing, genomics, and ocular amyloid (4). Subjects were followed longitudinally for up to 3 years.

Assessment of brain atrophy using MRI is a widely accepted measurement of disease progression in neurodegenerative disease (6, 7). However, the non-linearity of the magnetic field gradients produced by MRI scanners can produce errors in volumetric measurements leading to increased variability (8–10). This inconsistency reduces statistical power to detect subtle group differences or follow the progression of diseases in longitudinal and multi-site studies such as ONDRI (11, 12). Characterization and correction of scanner specific image distortions is an important requirement to assess neuroimaging biomarkers in large studies involving multiple sites with MRI scanners from different vendors (7, 11, 12).

System dependent geometric distortions in MRI may arise from imperfections in scanner hardware that cause magnetic field inhomogeneities (13, 14) and gradient field non-linearities (15, 16). However, gradient field non-linearities are the most prominent source of geometric distortions in structural MRI (11, 12, 17). Modern MRI scanners do incorporate 2D or 3D correction algorithms for such distortions (11, 18). However, all scanners do not enable them by default and varying distortion correction methods may be used across different scanners in multi-site studies. Moreover, some distortions may still present in images even after applying vendor supplied correction algorithms (19). Several groups have proposed post-processing algorithms to estimate and correct geometric distortions using 3D phantom scans (11, 15, 20, 21). There are two main techniques reported in the literature: (a) direct mapping of geometric distortions in each gradient field using specialized hardware to calculate the spherical harmonic expansion that can be used to correct the distortions (11, 21), and (b) indirect estimation of deformation fields using a phantom with identifiable structures (15, 20). The Alzheimer’s Disease Neuroimaging Initiative (ADNI) study employed the latter method using a custom-built phantom to be scanned *each time* a subject was scanned to estimate geometric distortion fields for subsequent correction of T1-weighted images and voxel scaling (22). The ONDRI study used a geometric phantom made out of Lego DUPLO^®^ bricks (12, 20) to estimate and correct gradient field induced geometric distortions at each imaging site.

In the ONDRI study, each imaging site acquired monthly 3D T1-weighted MR images of the geometric phantom to quantify geometric distortions (23). The objective of the current study was to measure the magnitude of the geometric distortions across sites, and to develop a distortion correction approach that minimized the variance of brain volumetrics. We defined a metric to quantify the geometric distortion from the measured distortion fields obtained from each scan session. Geometric distortions were compared across imaging sites and longitudinally over a 12-month period. The impact of gradient field induced geometric distortions on the volumetric measurements of specific structures was assessed using synthetic and human *in vivo* 3D T1-weighted MR images. We hypothesized that the correction of geometric distortions using a geometric phantom would lead to increased volumetric measurement precision associated with this multi-site longitudinal study.

## Methods

The neuroimaging protocol of the ONDRI study is designed to measure brain anatomy, microstructure, and resting-state functional networks in neurodegenerative disease cohorts using an MRI protocol that was previously described (23), which is consistent with the Canadian Dementia Imaging Protocol (CDIP) (24). The imaging protocol was designed to provide consistency between vendors. Briefly, each MRI session included the following scans: 3D T1-weighted anatomical, proton density (PD)/T2-weighted (PD/T2), fluid-attenuated inversion recovery (FLAIR), gradient echo, resting state functional MRI, and diffusion tensor imaging.

ONDRI participants were scanned at multiple time points at ten imaging sites across Ontario (4) incorporating three different MRI vendors: one Siemens (Erlangen, Germany) Prisma Fit, two Siemens Tim Trio systems, one Siemens Skyra, one General Electric (GE, USA) Signa HDxt, three GE MR750 scanners, and two Philips (Netherlands) Achieva systems. Each MRI site was required to acquire monthly scans of the MRI geometrical phantom built using Lego DUPLO^®^ bricks (12, 20) to estimate and correct geometric distortions due to gradient field non-linearities. Every site had their own LEGO phantom constructed to the same specifications. The manufacturer supplied built-in distortion corrections were also enabled during both human and phantom scans.

### Phantom Data Acquisition Protocol

Two T1-weighted scans of the MRI geometrical phantom were acquired in each monthly phantom scan session using the same MRI scan protocol as that used for the ONDRI 3D T1-weighted human scans. Details of the imaging protocol vary on each scanner and are provided in Scott *et al*. (23). The main difference for the phantom scan was that the FOV was increased in the sagittal direction by increasing the number of slices to 224 (from 176 or 180) to ensure full coverage of the phantom without aliasing. Different head coils were used in some ON-DRI imaging sites to accommodate the slightly larger size of the phantom as the coil used does not have a major effect on the geometric distortions. The first scan was acquired with the phantom positioned at the magnet isocentre. Then the bed was moved 5 cm into the scanner for a second scan. The shifted scan was used to increase the coverage of the phantom in the axial directions (20). The two phantom scans were registered to the known template of the MRI geometrical phantom to calculate the nonlinear distortion transformation needed to correct geometric distortions present within *in vivo* 3D T1-weighted scans as described previously (12, 20). Both automatic and manual quality assurance procedures were performed on each set of phantom images to ensure images were free of artefacts and acquired using the proper parameters (23).

### Quantification of Geometric Distortions

Geometric distortions were quantified using the distortion fields estimated for each of the MRI geometrical phantom scans. For example, a point, *p*(*x, y, z*) defined on an uncorrected scan is moved to a location defined by the point, *p′* (*x′, y′, z′*) in the corrected images (Figure 1). The magnitude of the distortion (*d*) at each point *p′*(*x′, y′, z′*) in the corrected image is then defined by the Euclidean distance given in Equation 1.

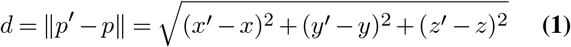

**Fig. 1.**
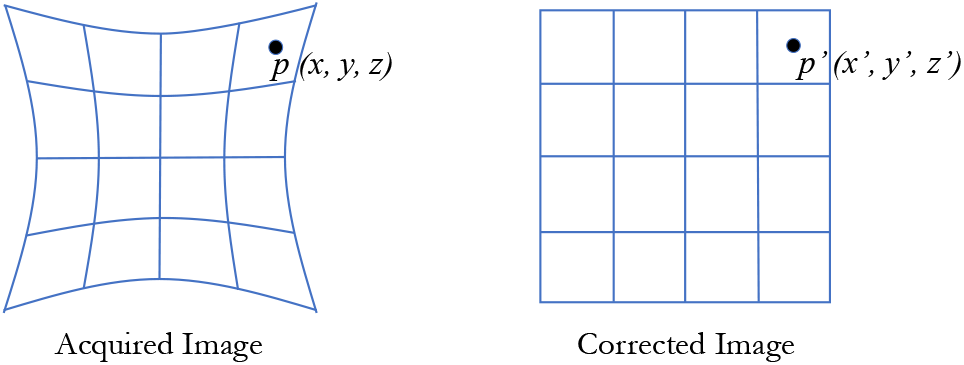
Geometric distortion correction: a point *p*(*x, y, z*) in the acquired image with geometric distortions is moved to *p′*(*x′, y′, z′*) in the corrected image.

The distortion metric (*d*_*m*_) was defined as the mean of the distortion (*d*) measured at all points distributed isotropically at 1 mm intervals within a spherical region of radius 100 mm centered at the magnet isocenter. The scalar value *d*_*m*_ defines the magnitude of the distortion estimated from the associated phantom scan for a specific time point at each of the ONDRI imaging sites. The MRI geometrical phantom scan sessions from 12 consecutive months were selected from each ONDRI imaging center to assess longitudinal variations in the geometric distortions at individual imaging sites. The distribution of the mean distortion, *d*_*m*_ was also compared between different imaging sites.

### Calculation of Volumetric Measurement Variability

#### Synthetic Images

A simulated cube of dimensions (2 × 2 × 2) cm^3^ was moved along *x, y* and *z* directions from −15 cm to 15 cm in 1 cm steps to assess the magnitude of volumetric measurement error at each site. The simulated distortion corrected volume was calculated at every time point for each imaging site. Variations across imaging sites were compared to evaluate the effects of geometric distortions on the volumetric measurement.

#### Quantitative MRI (qMRI) phantom

The CaliberMRI system standard quantitative MRI (qMRI) phantom (25) was modeled after the phantom designed for the ADNI MRI core (3). The qMRI phantom consists of a deionized water-filled spherical shell of 200 mm inner diameter. Inside the spherical shell, there is a frame consisting of 5 plates rigidly connected with positioning rods. The plates support 56 fiducial spheres, a 14-element T1 array, a 14-element T2 array, a 14-element proton density (PD) array, 2 resolution insets, and 2 wedges for slice profiling (schematic diagram in Figure 2a). T1, T2 and PD array elements were filled with varying concentrations of *NaCl*_2_, *MnCl*_2_, and *D*_2_*Cl* solutions, respectively to produce a gradual increase in MRI signal intensity from element 1 to 14 in each array (Figure 2b). The same qMRI phantom was scanned at ONDRI imaging sites as part of a larger travelling human subject phantom study examining intra- and inter-site scanner variance in the Ontario Brain Institute’s Integrated Discovery Projects (26). Each scanning session was comprised of a high-resolution 3D T1-weighted and a PD/T2-weighted pulse sequence as used in the ONDRI study.

**Fig. 2.**
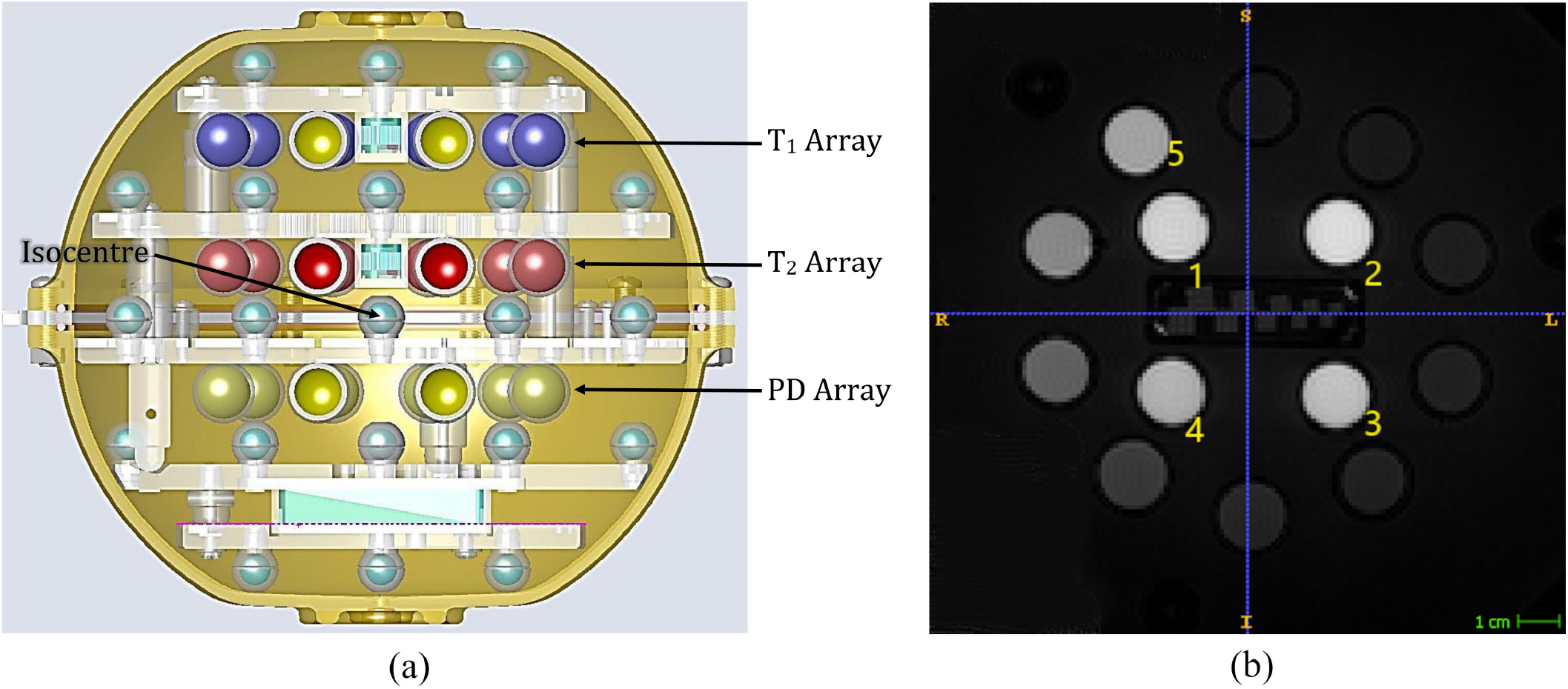
The HPD (CaliberMRI) system standard quantitative MRI (qMRI) phantom (25). (a) Cross-sectional view and (b) Coronal view of T1W scan showing T2 spheres. Figure 2a is reproduced with permission from CaliberMRI, a subsidiary company of High Precision Devices in Boulder, Colorado, United States (https://qmri.com/).

Geometric distortions in each qMRI phantom scan were corrected by applying the inverse of distortion field generated using the geometric phantom scans corresponding to the median distortion metric, *d*_*m*_ in the preceding six months at each respective ONDRI imaging site. The five highest intensity elements from the PD, T1, and T2 arrays were selected for volumetric analysis. Only the five highest intensity elements of each array were selected to avoid segmentation errors due to low contrast edges. All 15 spheres were segmented in the 3D T1-weighted images using semi-automatic tools; specifically, using clustering-based initialization followed by “evolution” using ITK-SNAP (27). The same user segmented the selected spheres using identical parameters in acquired and distortion-corrected MRI scans.

The variability of volumetric measurements was estimated by the standard deviation of the volume distribution of each sphere measured across the ONDRI imaging sites. Distance measurements were calculated between corresponding spheres in the PD and T2 arrays, and between the T2 and T1 arrays. The standard deviation of the distribution of each distance measurement across all sites was used to estimate the variability of distance measurements. Uncorrected and corrected measurement variabilities were compared using unpaired *t*-test (*p <* 0.05 was considered statistically significant).

Sample size estimates were performed to determine the minimum samples sizes to detect a 1%, 5%, and 10% change in volume before and after geometric correction as described in (28). For these calculations, *α* = 0.05 for a two tailed comparison, and power was set to 0.80. An initial volume of 3000 mm^3^ was used to represent a small brain structure. The average standard deviation of the volume distribution of each sphere measured across the ONDRI imaging sites before and after correction were used to estimate the variance.

#### Travelling Human Subjects

Two normal male subjects aged between 40-55 years were scanned at ONDRI imaging sites as a part of a travelling human subject study examining intra- and inter-site scanner variance in the Ontario Brain Institute’s Integrated Discovery Projects (26). Each scanning session included acquisition of high-resolution 3D T1-weighted and PD/T2-weighted images as in the ONDRI study. Geometric distortions in each 3D T1-weighted scan were corrected by applying the distortion field generated using geometric phantom scans corresponding to the median distortion metric, *d*_*m*_ in the preceding six months at each respective ONDRI imaging site. Three global tissue types; Grey Matter (GM), White Matter (WM), and Cerebrospinal Fluid (CSF) were segmented in the 3D T1-weighted scans using FMRIB’s Automated Segmentation Tool (FAST) using default settings (29) in FMRIB Software Library (FSL) (30). Brain extractions used in FAST were produced by the Brain Extraction Tool (BET) (31) in FSL using a fractional intensity threshold of 0.4 and the “robust” brain centre estimation option to minimize variability in the segmentation process.

For each of the two subjects, the coefficient of variance (CV) of the distribution of each tissue volume measured across the ONDRI imaging sites before and after correction was used to estimate the variance of the volumetric measurements. The CV of the uncorrected and corrected tissue volumes were statistically compared using two-way ANOVA (*p <* 0.05 considered statistically significant).

#### Cross-Sectional Measures of Human Brain Structures

Base-line imaging data from 150 people with cerebrovascaular disease (CVD) acquired at nine ONDRI imaging sites were selected to assess the effects of geometric distortion corrections on MRI-derived volumetric measurements of the human brain. Brain volumetry was performed using a semi-automatic pipeline called Lesion Explorer (LE) (32, 33) as published previously. The volumes of the following six different global tissue and lesion classes were measured to assess the magnitude of changes following geometric distortion corrections: normal appearing white matter (NAWM), normal appearing grey matter (NAGM), sulcal cerebrospinal fluid (sCSF), ventricular cerebrospinal fluid (vCSF), periventricular white matter hyperintensities (pWMH), and deep white matter hyperintensities (dWMH). For each subject’s scan, the appropriate geometric distortion correction was applied to the 3D label map containing all six regions using the distortion field calculated from the MRI of the geometrical phantom. Nearest-neighbor interpolation was used during the image warping operations to keep labels intact. Since the geometric distortion correction field was not obtained at the time of imaging each subject, several different correction strategies were attempted to determine the method that minimized the variability of the volumetric measurements. The following three approaches were compared: 1) using the phantom scan session that corresponded to the maximum *d*_*m*_ of the selected period for geometric distortion analysis, 2) using the phantom scan session that corresponded to the median *d*_*m*_ of the selected period, and 3) using the phantom scan session that corresponded to the minimum *d*_*m*_ of the selected period. The volumetric measurements of the acquired and corrected scans for each selected tissue/lesion class were statistically compared using repeated measures ANOVA followed by Tukey’s multiple comparisons test (for normally distributed data) and Friedman test followed by Dunn’s multiple comparison test (for non-parametric data). In all cases *p <* 0.05 was considered statistically significant.

## Results

### Quantification of Geometric Distortions

Geometric distortion fields were generated from monthly phantom scans at ONDRI imaging sites for all phantom scans that passed quality assurance checks. A total of 116 valid scans were available from 10 ONDRI imaging sites. One site had only 10 valid scans and two other sites had 11 scans while the remaining seven sites had all 12 scans. The mean distortion metric, *d*_*m*_ for each geometric field was estimated. Figure 3a shows the time variation of *d*_*m*_ at every imaging site. Except for a few outliers, geometric distortions within a site remained similar throughout the 12-month period. Figure 3b shows the average and standard deviation of *d*_*m*_ for all times points at each ONDRI imaging site. Figure 4 shows example phantom scans from sites with notably different geometric distortions and the corresponding corrected images. The average, median, minimum, and maximum *d*_*m*_ are given in Table 1 for all ONDRI sites. The magnitude and reproducibility of geometric distortion varied considerably at different sites.

**Table 1.**
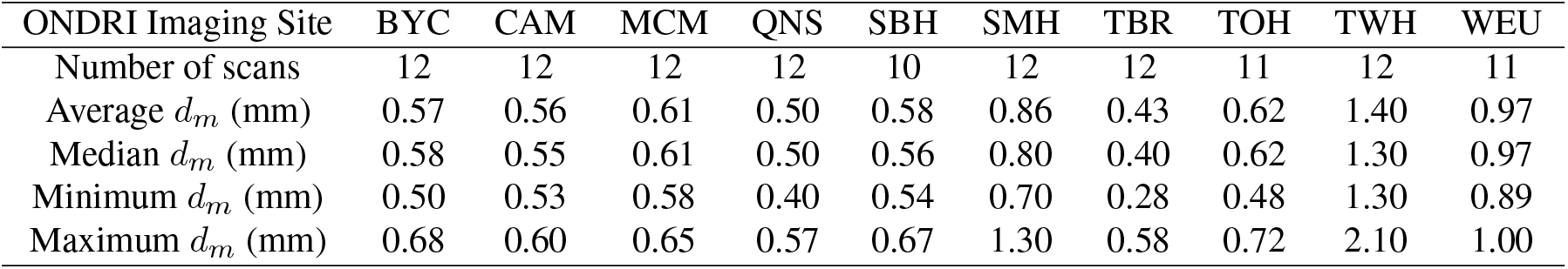
Geometric distortion metric, *d*_*m*_, estimated using monthly phantom scans at ONDRI imaging sites. BYC = Baycrest Centre (Siemens 3T Tim Trio), CAM = Centre for Addictions and Mental Health (GE 3T MR750), MCM = McMaster St. Joseph’s Hospital (GE 3T MR750), QNS = Queen’s University (Siemens 3T Tim Trio), SBH = Sunnybrook Hospital (GE 3T MR750), SMH = St Michael’s Hospital (Siemens 3T Skyra), TBR = Thunder Bay Research Institute (Philips 3T Achieva), TOH = Ottawa Hospital (Siemens 3T Tim Trio), TWH = Toronto Western Hospital (GE 3T Signa HDxt), WEU = Western University (Siemens 3T Prisma).

**Fig. 3.**
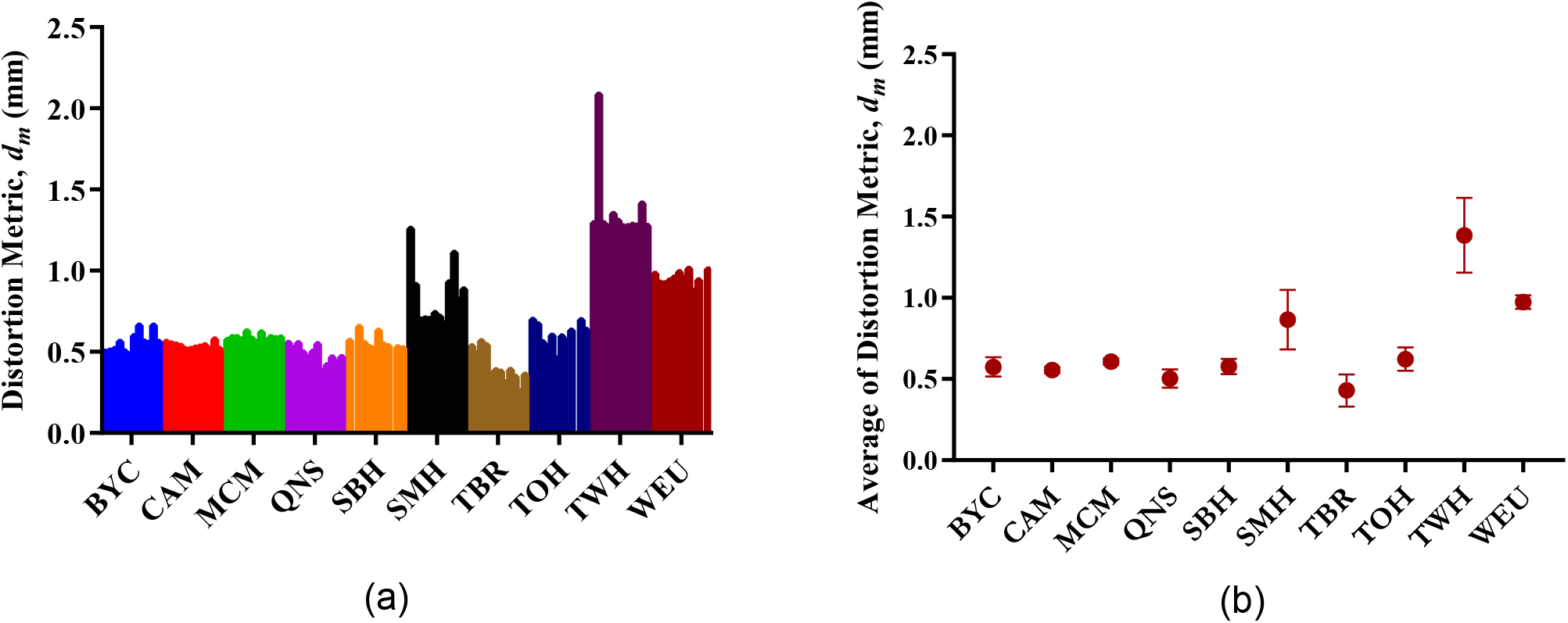
(a) Monthly distortion metric, *dm* value across the 12-month period and (b) the average of *dm* at each ONDRI imaging site. BYC = Baycrest Centre (Siemens 3T Tim Trio), CAM = Centre for Addictions and Mental Health (GE 3T MR750), MCM = McMaster St. Joseph’s Hospital (GE 3T MR750), QNS = Queen’s University (Siemens 3T Tim Trio), SBH = Sunnybrook Hospital (GE 3T MR750), SMH = St Michael’s Hospital (Siemens 3T Skyra), TBR = Thunder Bay Research Institute (Philips 3T Achieva), TOH = Ottawa Hospital (Siemens 3T Tim Trio), TWH = Toronto Western Hospital (GE 3T Signa HDxt), WEU = Western University (Siemens 3T Prisma).

**Fig. 4.**
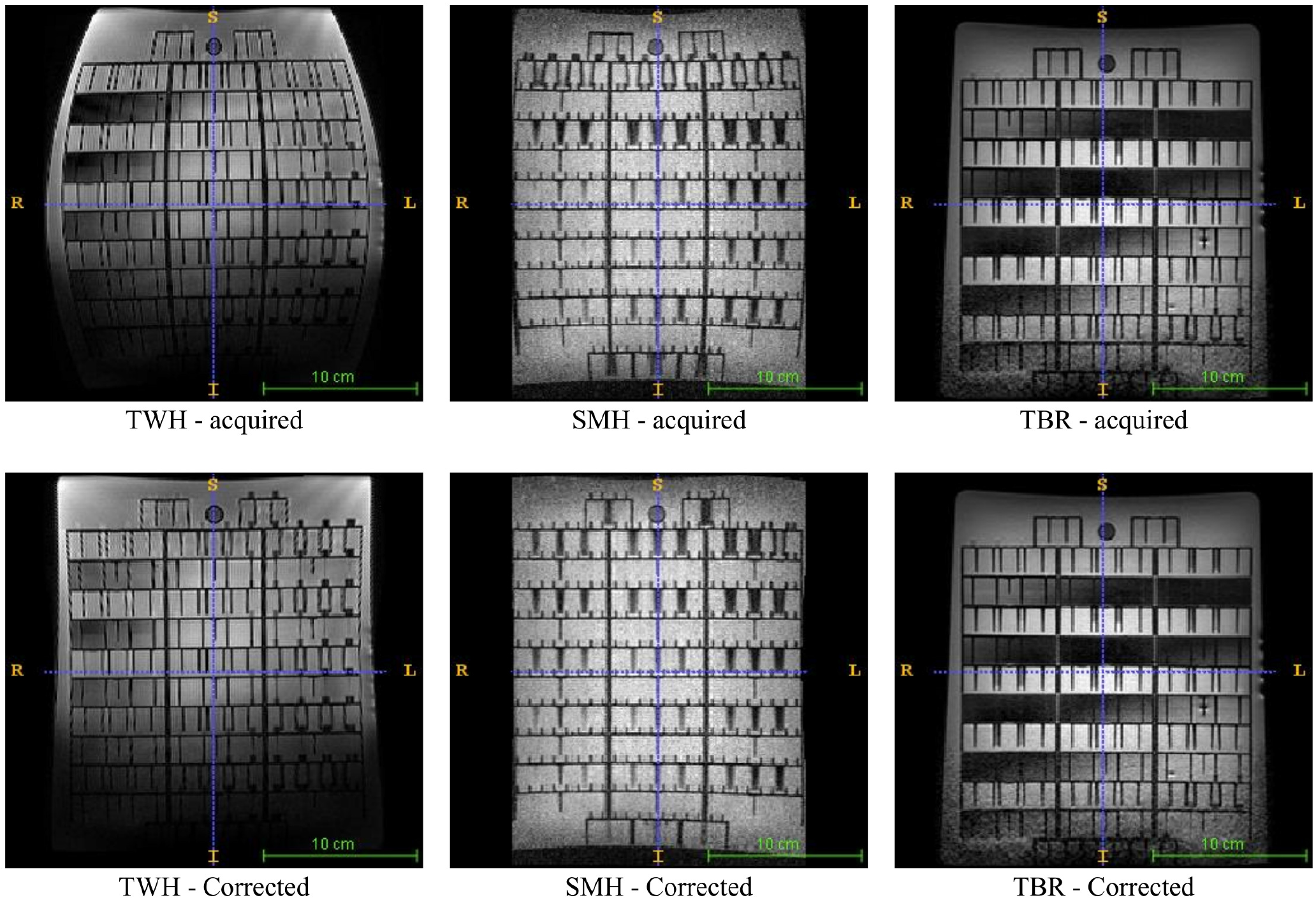
Phantom scans before (top) and after (bottom) distortion correction from three different sites. TWH = Toronto Western Hospital (GE 3T Signa HDxt), SMH = St Michael’s Hospital (Siemens 3T Skyra), TBR = Thunder Bay Research Institute (Philips 3T Achieva).

### Volumetric Measurement Variability on Synthetic Images

The uncorrected volume of a simulated 8 cm^3^ cube as it is moved along *x, y* and *z* axes is shown in Figure 5. These curves represent the effect of the average geometric distortion across 12 months. The origin (0, 0, 0) of the coordinate corresponds to the magnet isocenter. The volumetric changes increased as the distance to magnet isocenter increased in all three axes, but the magnitude varied between axes. Distortions caused increased or decreased volumes independently along the three directions. Some sites also showed much greater distortions than others, particularly in the *x* and *z* axes. These data suggest that the geometric distortions were non-linear with different magnitudes along the three gradient directions.

**Fig. 5.**
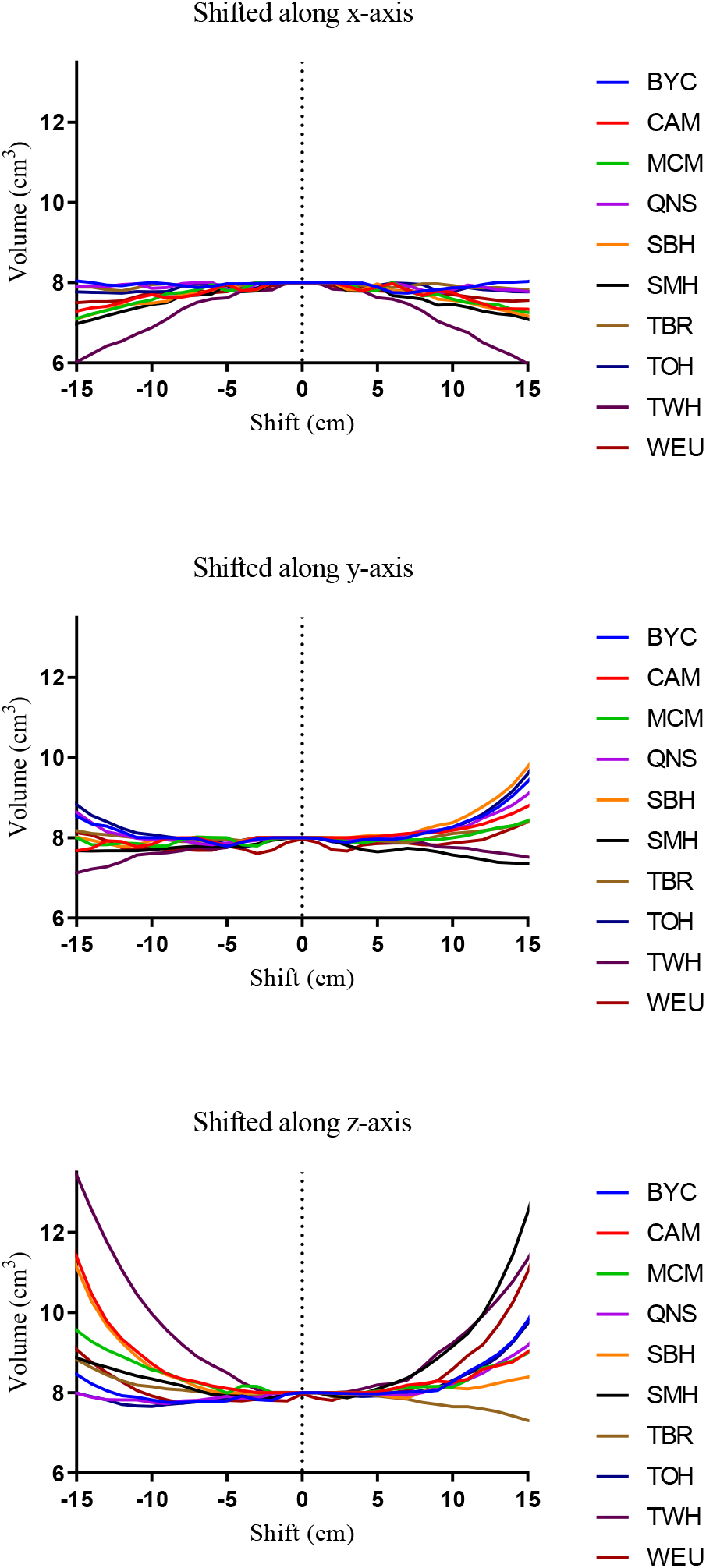
Average volume of all time points of geometric distortion corrected simulated cube with dimensions (2 *×* 2 *×* 2) cm^3^ at ONDRI imaging sites. BYC = Baycrest Centre (Siemens 3T Tim Trio), CAM = Centre for Addictions and Mental Health (GE 3T MR750), MCM = McMaster St. Joseph’s Hospital (GE 3T MR750), QNS = Queen’s University (Siemens 3T Tim Trio), SBH = Sunnybrook Hospital (GE 3T MR750), SMH = St Michael’s Hospital (Siemens 3T Skyra), TBR = Thunder Bay Research Institute (Philips 3T Achieva), TOH = Ottawa Hospital (Siemens 3T Tim Trio), TWH = Toronto Western Hospital (GE 3T Signa HDxt), WEU = Western University (Siemens 3T Prisma).

### Analysis of measurement variability using a quantitative MRI (qMRI) phantom

Measurements of the volume of each sphere within the qMRI phantom should produce the same value at each imaging site since it is the same physical object. Figures 6a and 6b show the distribution of all volumetric measurements of selected spheres across all ON-DRI imaging sites before and after gradient distortion correction. A lower variability in measurement distributions is evident after geometric distortion corrections, where the average standard deviation reduced to 125 mm^3^ after correction from 192 mm^3^ (35% reduction). The measurement variability associated with each sphere across ONDRI imaging sites was estimated by the standard deviation (SD) of the volumetric measurement distribution before and after correction. The SD of these volumes before and after correction, provided in Figure 6c, were normally distributed and showed a statistically significant (*p <* 0.01) mean difference of −67 ± 20 mm^3^ (Figure 6d).

**Fig. 6.**
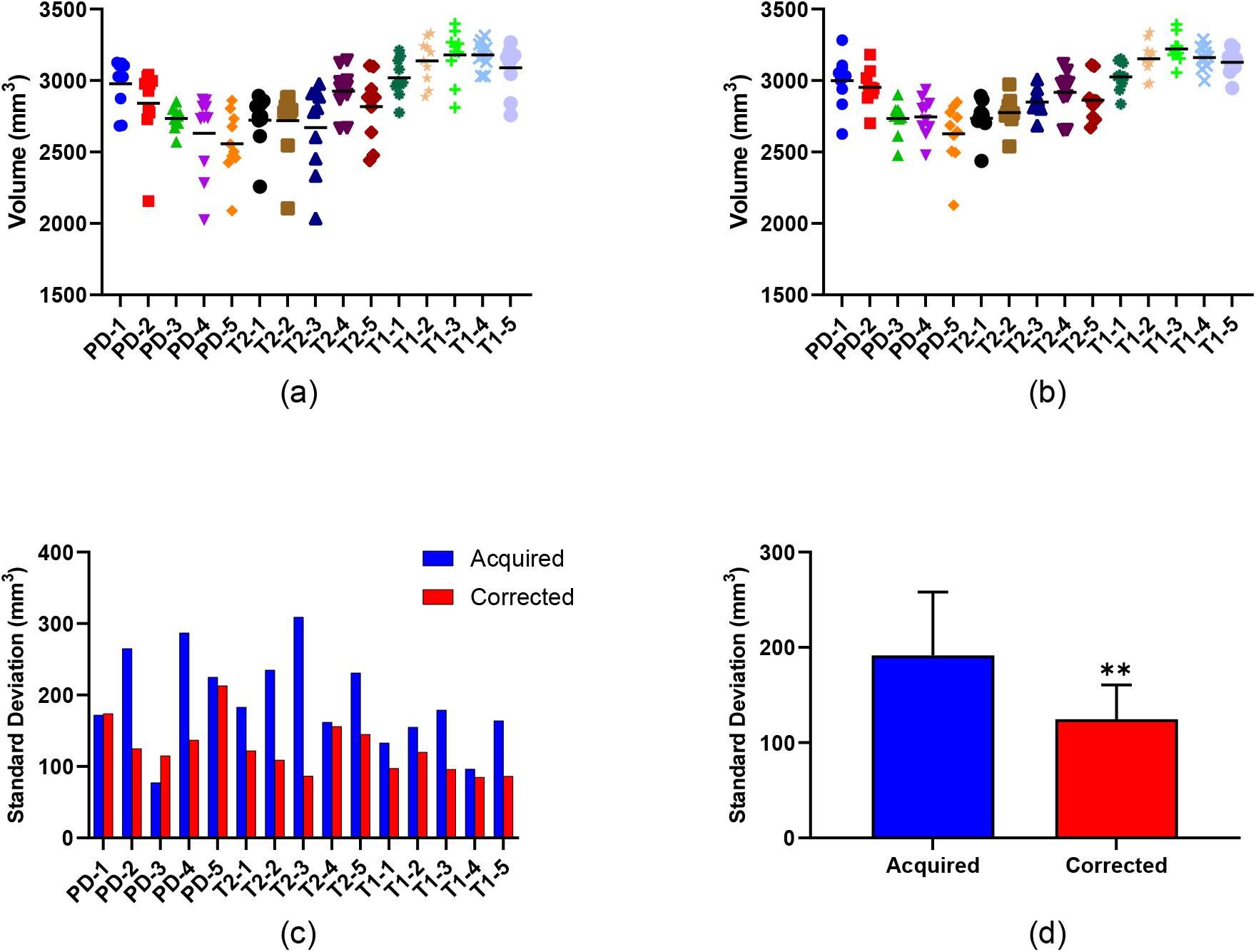
The volumes of each selected sphere measured on the (a) acquired and (b) corrected T1-weighted scans at each ONDRI imaging site and corresponding (c) the distribution and (d) the mean of the standard deviations (SD) (** indicates a statistically significant difference with *p <* 0.01). PD-1 to PD-5 = five highest intensity spheres of the PD array, T2-1 to T2-5 = five highest intensity spheres of the T2 array, and T1-1 to T1-5 = five highest intensity spheres of the T1 array.

The variability associated with linear distances between corresponding selected spheres in the PD and T2 sphere arrays, and between T2 and T1 sphere arrays in scans at all imaging sites were also compared (Figures 7a and 7b show the distribution of all distance measurements across all ONDRI imaging sites). A lower variability in measurement distributions is evident after geometric distortion corrections, where the average standard deviation reduced to 0.08 mm after correction from 0.14 mm (43% reduction). The measurement variability associated with the distance between spheres across ONDRI imaging sites was estimated using the standard deviation (SD) of the distance measurement before and after correction. The SD of this distance measurement before and after correction (Figure 7c), were normally distributed and showed a statistically significant (*p <* 0.001) mean difference of −0.062 ± 0.009 mm (Figure 7d).

**Fig. 7.**
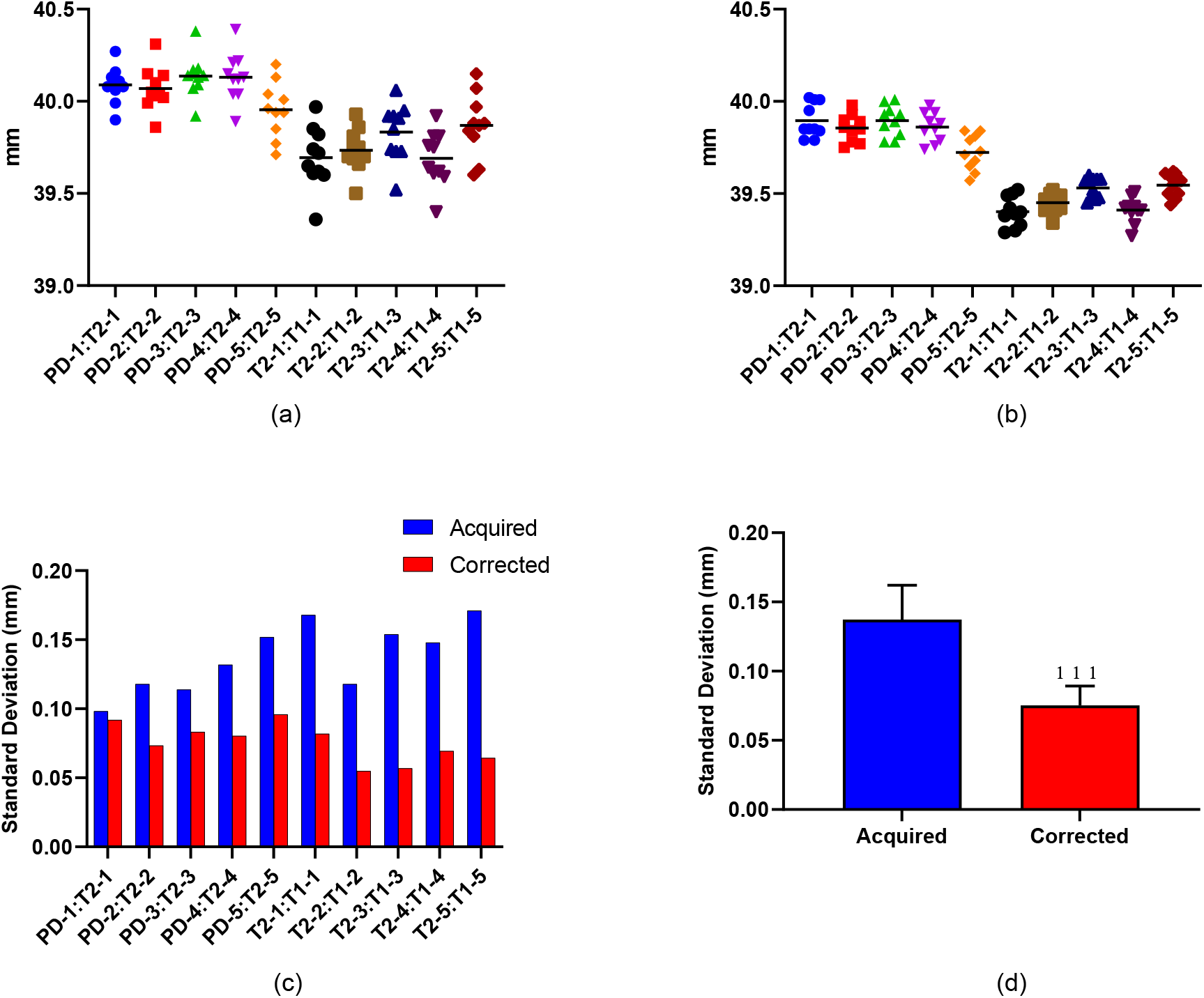
The distance measurements between corresponding selected spheres in the PD and T2 arrays, and between the T2 and T1 arrays on the (a) acquired and (b) corrected T1-weighted scans at each ONDRI imaging site and corresponding (c) the distribution and (d) the mean of the standard deviations (SD) (** indicates a statistically significant difference with *p <* 0.01). PD-1 to PD-5 = five highest intensity spheres of the PD array, T2-1 to T2-5 = five highest intensity spheres of the T2 array, and T1-1 to T1-5 = five highest intensity spheres of the T1 array.

The measurement variability in volumetric measurements decreased significantly. This reduction of measurement variability would improve the power to detect differences in volumes between disease cohorts or would decrease the number of participants needed for clinical trials to show a statistically significant effect. Sample size estimates (two-tailed, *α* = 0.05, power = 0.8) demonstrate that fewer subjects are needed to observe significant differences between groups. To detect a 1%, 5% or 10% change in volume would have required 643, 26, and 7 subjects per group, respectively before correction, which reduces to 273, 11, and 3 subjects per group, respectively after correction.

### Measurements Variability in Travelling Human Subjects

The volume of each tissue type should ideally produce the same value for a subject at all imaging sites. However, there is a variability in measured volumes due to multiple factors including MRI scanner-introduced geometric distortions at different sites. The measurement variability associated with each tissue type across ONDRI imaging sites estimated by the Coefficient of Variance (CV) of the volumetric measurement distribution before and after correction for both subjects are provided in Figure 8. The variance of volumes across ONDRI imaging sites estimated by CV was reduced in every tissue type of both subject scans. Results of the statistical comparison of changes of CV using two-way ANOVA showed a statistically significant (*p <* 0.01) reduction of variance in corrected volumetric data across ONDRI imaging sites.

**Fig. 8.**
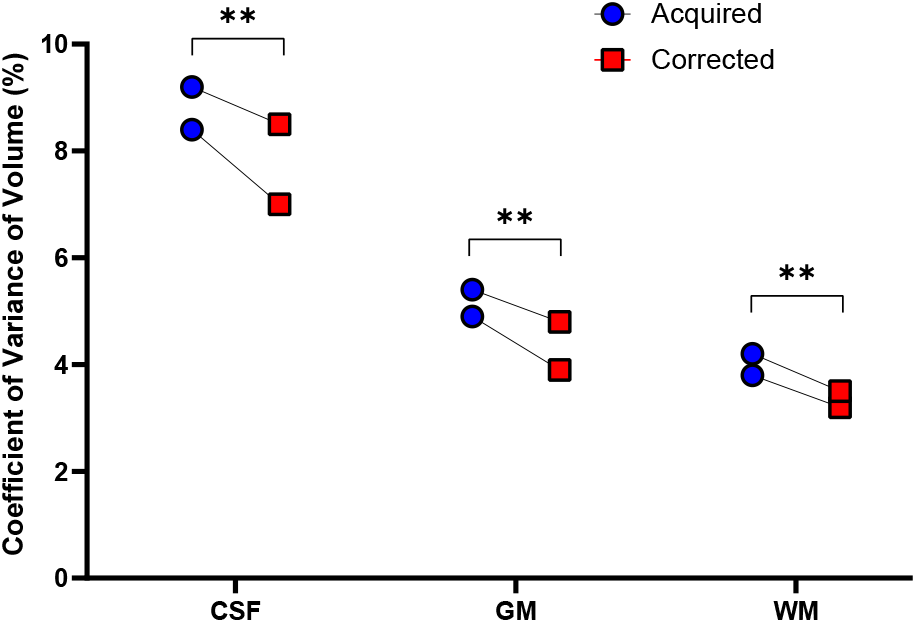
The coefficient of variance (CV) changes of each selected tissue type measured for both subjects on acquired and corrected T1-weighted scans at 10 ONDRI imaging sites (** indicates a statistically significant change of *p <* 0.01). A significant decrease in CV was found with two-way ANOVA due to the applied geometric distortion correction.

### Variability in the Volume of Human Brain Structures

The volumetric data for six selected tissue types were first tested for normality using both the Shapiro-Wilk test and the Kolmogorov-Smirnov test. The volumes of NAWM (*p* = 0.52 and *p >* 0.10), and NAGM (*p* = 0.92 and *p >* 0.10) were normally distributed while the volumes of sCSF (*p* = 0.04 and *p* = 0.02), vCSF (*p <* 0.001 for both tests), pWMH (*p <* 0.001 for both tests), and dWMH (*p <* 0.001 for both tests) were non-parametric. Table 2 provides the mean volume of the tissue types before and after each correction. Statistical comparisons were performed according to the nature of the distribution of each tissue volume. The results of these comparisons are summarized in Table 3 (volume or rank differences for each subject is shown in Figure 9). Incorporating a geometric distortion correction using the maximum, median, or minimum distortion field acquired during the selected 12-month period always produced a statistically significant volume decrease compared to the uncorrected data. The average magnitude of this decrease was ∼5 cm^3^ or 1.3% in NAWM and ∼9 cm^3^ or 1.7% in NAGM (Table 3). However, volume differences between corrected volumes using the maximum or minimum distortion correction fields acquired at different times during the 12-month period were not always statistically different (Table 3).

**Table 2.** Mean volume of all tissue types in people with cerebrovascular diseases before (Acquired) and after geometric distortion correction using geometric distortion fields corresponding to the maximum, median, and minimum distortion metric, *d*_*m*_.

| | Mean Volume ± Standard Deviation (cm<sup>3</sup>) | | | |
| --- | --- | --- | --- | --- |
|  | Acquired | Maximum | Median | Minimum |
| NAWM | 387.1 ± 55.3 | 381.6 ± 53.9 | 381.8 ± 54.0 | 382.6 ± 54.1 |
| NAGM | 536.1 ± 53.2 | 527.3 ± 52.3 | 527.0 ± 52.0 | 528.1 ± 51.8 |
| sCSF | 243.3 ± 61.2 | 239.3 ± 59.8 | 239.1 ± 59.8 | 239.8 ± 60.3 |
| vCSF | 41.4 ± 23.2 | 41.1 ± 23.0 | 41.1 ± 23.0 | 41.2 ± 23.1 |
| pWMH | 9.01 ± 12.38 | 8.92 ± 12.25 | 8.92 ± 12.24 | 8.94 ± 12.26 |
| dWMH | 0.96 ± 1.21 | 0.95 ± 1.19 | 0.96 ± 1.20 | 0.96 ± 1.19 |

**Table 3.** Average differences in measured tissue volumes before and after gradient distortion correction and differences between gradient distortion corrections corresponding to the maximum, median, and minimum distortion metric, *d*_*m*_. The mean difference is reported for normally distributed data compared with Tukey’s multiple comparisons test and the rank sum difference is reported for non-parametric data compared with Dunn’s multiple comparison test. Statistically significant values with *p <* 0.05 are marked with ^*\**^, and *p <* 0.001 are marked with ^*\*\*\**^.

|  | Mean Difference (cm<sup>3</sup>) |  | Rank Sum Difference |  |  |  |
| --- | --- | --- | --- | --- | --- | --- |
|  | NAWM | NAGM | sCSF | vCSF | pWMH | dWMH |
| Acquired vs. Maximum | 5.52*** | 8.79*** | 266*** | 174*** | 185*** | 130*** |
| Acquired vs. Median | 5.33*** | 9.09*** | 322*** | 161*** | 173*** | 125*** |
| Acquired vs. Minimum | 4.51*** | 7.95*** | 261*** | 112*** | 125*** | 92*** |
| Maximum vs. Median | -0.19 | 0.30 | 56 | -13 | -13 | -5 |
| Maximum vs. Minimum | -1.01*** | -0.84* | -5 | -62* | -61* | -39 |
| Median vs. Minimum | -0.82*** | -1.14*** | -61* | -49 | -48 | -33 |

**Fig. 9.**
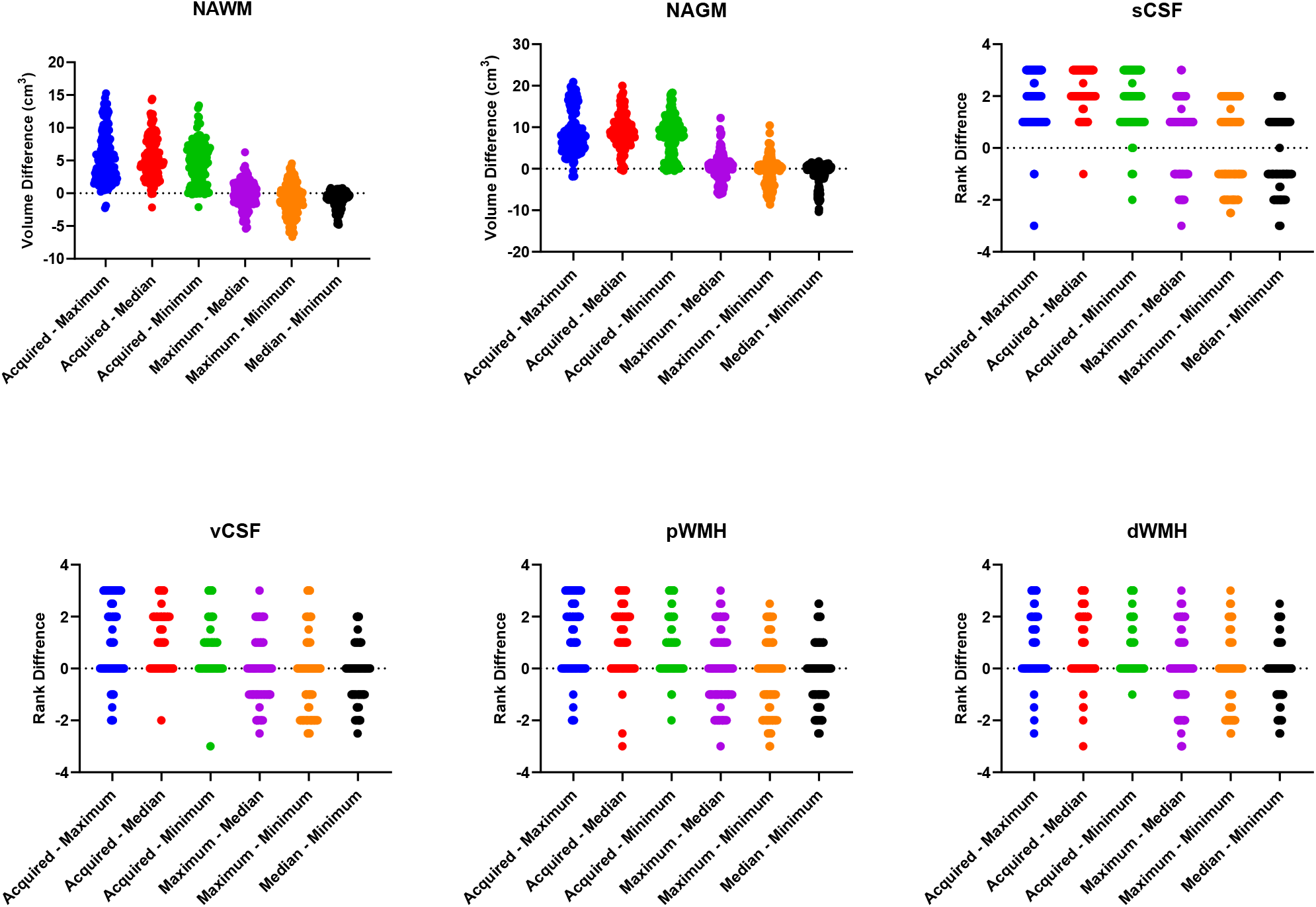
Tissue volume comparisons: The volume difference (cm^3^) for normally distributed data (NAWM and NAGM) was compared with repeated measures ANOVA, while rank difference for non-parametric data (sCSF, vCSF, pWMH, and dWMH) was compared with Friedman test.

## Discussion

The overall goal of this study was to determine whether the correction of geometric distortions in high resolution 3D T1-weighted MRI scans using monthly acquisitions of a geometric phantom to map deformation fields could increase the precision of brain volumetric measurements associated with multi-site studies. Geometric distortions were found to vary substantially between different MRI scanners (including scanners from different vendors), but were relatively stable on each scanner over a twelve-month interval. Geometric distortions were also found to vary spatially, increasing in severity with greater distance from the magnet isocenter and as a function of gradient axis. Using the CaliberMRI standardized phantom, geometric distortion correction produced a 35% decrease in the standard deviation of internal sphere volume measurements, and a 42% decrease in the standard deviation associated with distance measurements between spherical phantoms. This increase in precision led to a substantial decrease in the predicted sample sizes needed to detect volumetric changes. The increased measurement precision for brain volumetric data when incorporating geometric distortion correction across different scanners was confirmed using gray matter and white matter volume measurements of two travelling human subjects. The application of the distortion correction to human scans acquired across ten sites resulted in a small but statistically significant change in the volume of NAWM, NAGM, and sCSF compared to the uncorrected data.

This study estimated both volume change and linear displacement due to geometric distortion in MRI using a geometrical phantom. Within a spherical ROI of 100 mm radius centered at the magnet isocenter where distortions are minimal, the overall distortion metric, *d*_*m*_ developed to measure linear displacement, was found to vary between 0.43 mm and 1.39 mm for scanners at the ten ONDRI imaging sites in the 12-month period studied. Average volume difference of a simulated 8 cm^3^ cube moved along x, y and z axes in the same period varied between 0.2% and 28.3% across the ten scanners. These volume changes are very small at the isocenter, however both displacement and volume change increased substantially as the distance from the magnet isocenter increased.

A number of previous studies have estimated and corrected geometric distortions in MRI (11, 34–37), particularly in application areas such image-guided radiation therapy (17, 38) and image-guided surgery (39). Major studies of MRI geometric distortions on 1.5T and 3T scanners reported within last 15 years are summarized in Table 4. Most previous studies also compared geometric distortion between different scanners. When results were provided for various positions within the scanner, the distortions at locations closest to 100 mm away from the magnet isocenter were included in Table 4 for comparison to the current work. The geometric distortions in 7T MRI compared to 3T MRI reported by Lau et al. (39) were excluded as absolute distortion was not measured. Most previous studies did not report volume changes due to geometric distortion. Distortion values reported in the literature were estimated using the displacement of known points in a physical phantom. Mean distortion describes the average of displacements of all points used for the estimation, which differs from the distortion metric, *d*_*m*_ used in the current study. However, all distortion values quantify linear geometric distortions within a region of interest. As evident from Table 4, the measured distortions ranged from 0.23 mm to 3.2 mm, which is similar to the range of displacements measured across the scanners used at the ten sites included in the current study.

**Table 4.**
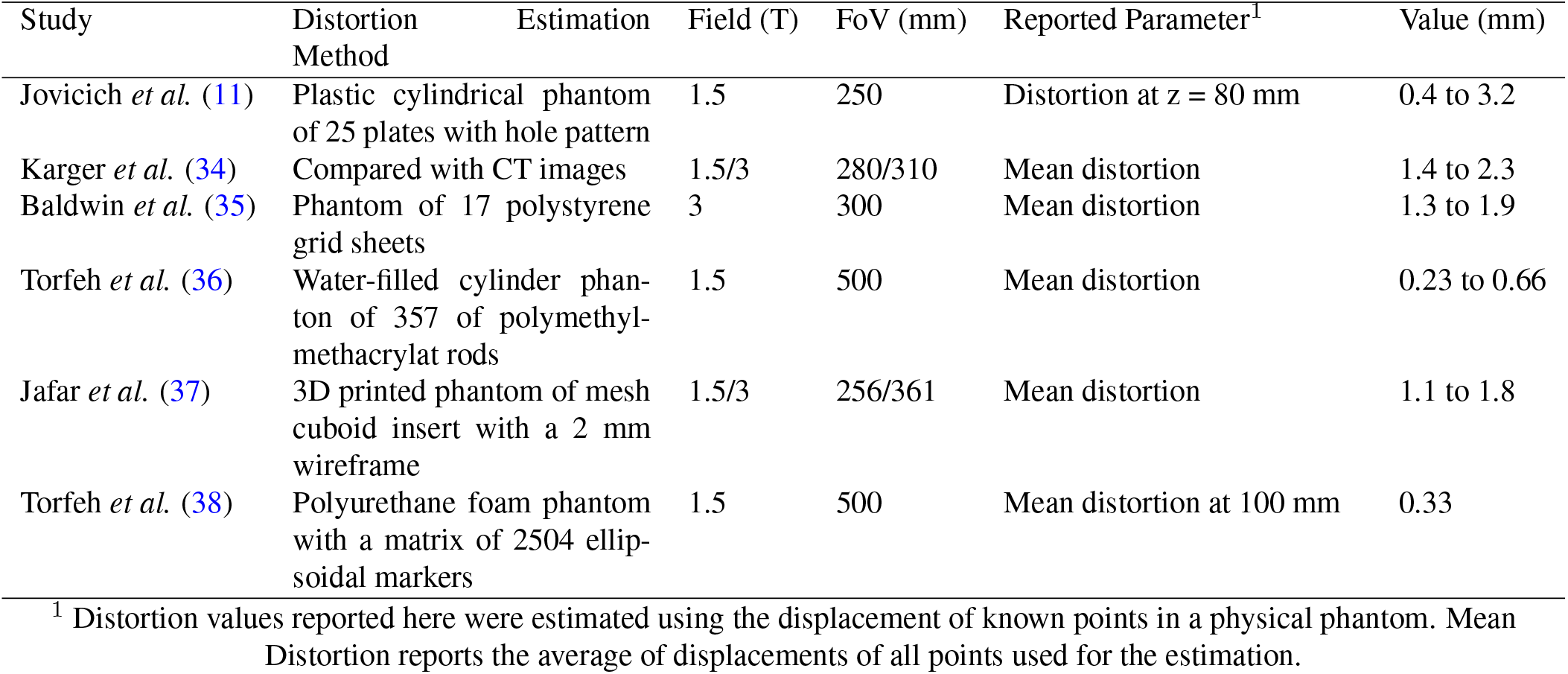
Summary of distortions reported in major studies of MRI geometric distortions

Imaging the same standardized (CaliberMRI) phantom containing high contrast spheres with a volume of ∼3000 mm^3^ at all imaging sites provided a unique opportunity to assess the impact of geometric distortion correction on volumetric measurements. A mean linear displacement between 0.20 and 0.32 mm and a mean sphere volume change between 1.4% and 6.4% across scanners was observed. Using measurements from this phantom, we estimated the potential reduction in samples sizes that could be achieved when measuring volumetric changes incorporating the gradient distortion correction. For example, to detect a 1% change in volume between groups, a substantial decrease in sample size could be achieved by incorporating gradient distortion correction to reduce variability between scanners. This result has important implications for multi-site imaging studies that incorporate volumetric outcome measures, whether related to the effects of drug treatment or natural history studies. The current study suggests that monthly phantom scans to track geometric distortions, and the use of these corrections, could significantly reduce the overall cost associated with such imaging studies, or boost statistical power. The data acquired in two participants that travelled to all ten imaging sites supports this result. The CVs associated with CSF, GM and WM measurements decreased by 11.9%, 15.5%, and 16.3% respectively from initial values following geometric distortion correction. While seemingly small, this increase in precision would lead to more efficient imaging trials at the expense of requiring regular phantoms scans and methods to apply geometric distortion corrections.

When examining the impact of distortion correction on *in vivo* average volume measurements of NAWH, NAGM and sCSF, it was found that these tissue volumes were over-estimated by 1.4%, 1.7% and 1.7%, respectively, without distortion correction. Volume measurements in other tissues remained virtually unchanged after distortion correction. Greater effects would be expected for regions near the periphery of the brain as geometric distortions are greater away from the magnet isocenter as shown in Figure 5. The volumes of structures near the magnet isocenter such as the ventricle (vCSF) are relatively unaffected.

There are several limitations to this work that should be considered. First, the distortion correction field estimated at the time of every human scan was not acquired and therefore could not be used to correct each image. Instead, this study examined whether monthly scans could improve measurement precision at a greatly reduced cost. The use of distortion corrections can also introduce uncertainties into the volumetric measurement process. For example, the calculation of the distortion field incorporates non-rigid registration of phantom scans with ideal structure (12, 20) which introduces some uncertainty. This uncertainty can be minimized to some degree by utilizing the median *d*_*m*_ within the six months preceding the scan for stable MRI systems. Another limitation of this work is that only a small subset of brain volumes was examined (e.g. NAWM, NAGM). Future studies may find larger effects in small volumes at the periphery of the brain. Finally, in the current study, all sites were advised to turn on built-in gradient distortion corrections provided on the scanners, but these instructions were not uniformly applied. Interestingly, most scanners had distortion corrections switched off by default. As a result, only a few sites used built-in 3D corrections, while others used only 2D correction or no correction. This result highlights both the potential benefit of using an independent geometric distortion protocol, and the need to routinely check whether sites are applying built in distortion correction methods.

## Conclusions

The current study demonstrates a significant reduction in the variance associated with multi-site volumetric measurements of a standardized phantom and human volunteers when incorporating a three-dimensional geometric distortion correction. Without geometric distortion correction, global measurements of normal appearing white matter, normal appearing gray matter and sulcal cerebrospinal fluid were overestimated by 1.4%, 1.7% and 1.7%, respectively in 150 people with cerebrovascular disease. Although the volume changes were small, the CaliberMRI standardized phantom analysis showed that a 35% decrease in the standard deviation of internal sphere volume measurements could be realized by incorporating distortion correction based on monthly scanner measurements. Based on these results, geometric distortion correction using monthly scans appears to be a cost-effective approach to increase the precision of volumetric MRI measurements in multi-site imaging studies. It may also be a safeguard against the inconsistent use of 2D and 3D built-in scanner distortion corrections in multi-site studies.

## ACKNOWLEDGEMENTS

This research was conducted with support from the Ontario Neurodegenerative Disease Research Initiative (ONDRI) through the Ontario Brain Institute, an independent non-profit corporation, funded partially by the Ontario government.

The authors would like to thank all ONDRI participants including the ONDRI investigators and the ONDRI governing committees: executive committee; steering committee; publication committee; recruiting clinicians; assessment platforms leaders; and the ONDRI project management team. For a full list of the ONDRI investigators, please visit: www.ondri.ca/people. The authors also acknowledge Tharushan Selliah, Abiramy Uthirakumaran, Ella Lawrence, and Aditi Chemparathy for their contributions to the Travelling Human Subjects Study.

